# Longitudinal Stability of Mood-Related Resting-State Networks in Youth with Symptomatic Bipolar-I/II Disorder

**DOI:** 10.1101/2025.01.10.630933

**Authors:** Danella M. Hafeman, Jamie Feldman, Jessica Mak, John Merranko, Tina R. Goldstein, Caterina Gratton, Mary L. Phillips, Boris Birmaher

**Affiliations:** University of Pittsburgh School of Medicine, Department of Psychiatry; Western Psychiatric Hospital, University of Pittsburgh Medical Center; Florida State University

**Keywords:** bipolar disorder, functional connectivity, fMRI, precision imaging, longitudinal studies, transition-age youth

## Abstract

Bipolar disorder (BD) is characterized by temporal instability of mood and energy, but the neural correlates of this instability are poorly understood. In previous cross-sectional studies, mood state in BD has been associated with differential functional connectivity (FC) amongst several subcortical regions and ventromedial prefrontal cortex. Here, we assess whether BD is associated with longitudinal instability within this mood-related network of interest (NOI). Young people with BD-I/II were scanned 4-6 times and healthy controls (HC) were scanned 4 times over 9 months. Following preprocessing of 20-minute resting-state scans, we assessed across-scan correlation of FC, focusing on FC between regions previously associated with BD mood state. Utilizing Bayesian models, we assessed the relationship between diagnostic group and within-person, across-scan correlation, adjusting for motion, time-of-day, and inter-scan interval; prediction intervals (PI) are reported. In a sample of 16 youth (11 BD, 5 HC; 16.3-23.3 years old) with 70 scans (50 BD, 20 HC), across-scan NOI stability was higher within-than between-person (0.70 vs. 0.54; p<.0001). BD (vs. HC) within-person scan-pairs showed lower NOI stability (mean -0.109; 95% PI -0.181, -0.038), distinguishing BD vs. HC with excellent accuracy (AUC=0.95). NOI instability was more pronounced with manic symptoms (mean -0.012; 95% PI -0.023, -0.0002) and in BD-II (vs. BD-I; mean -0.071; 90% PI -0.136, -0.007). Results persisted after accounting for medications, comorbidity, and sleep/arousal measures. Within this pilot sample, BD is characterized by less within-person stability of a mood-related NOI. While preliminary, these results highlight a possible role for precision imaging approaches to elucidate neural mechanisms underlying BD.

## INTRODUCTION

Bipolar disorder (BD) is a serious mental illness that, particularly if undiagnosed, is associated with impairment in psychosocial functioning, substance use, psychiatric hospitalizations, and suicidal thoughts and behaviors [1–3]. Peak onset of BD is in adolescence and young adulthood [4], when this disorder can also disrupt attainment of education, employment, and other milestones. Currently, average delays between initial mood episode and accurate diagnosis are over five years [5] and even longer in those with earlier age of onset [5, 6]. A neural marker with potential to identify youth with BD could have substantial clinical and public health implications. Studies utilizing functional MRI have pointed to reproducible differences in BD (vs. healthy controls; HC), particularly in regions such as the amygdala and striatum [7]; however, while important, these findings do not distinguish individuals with BD (vs. HC) on the individual level.

Given that BD is defined by episodic changes in mood and energy, it follows that the disorder may also be characterized by instability of brain networks that give rise to these mood changes. One way to assess brain networks is to evaluate the degree to which fluctuations in blood-oxygen level dependent (BOLD) signal are correlated across brain regions, even in the absence of stimuli [8]. This measure of correlation, known as functional connectivity (FC), has provided insights into the composition of brain networks and individual differences [9]. Recent developments in precision imaging methods have indicated that, given sufficient resting-state data (e.g., >20 minutes) and implementation of advanced denoising strategies, FC in healthy adults is relatively stable over time [10]. However, this network stability can be disrupted given perturbation, as shown by the acute and reversible effects of arm casting on the sensorimotor network [11]. Network stability may also be a marker of psychological functioning, higher in individuals with better social skills, and lower (and not showing appropriate developmental increases) in individuals with psychopathology [12, 13]. Given these recent and intriguing findings, it is possible that individuals with BD may also show longitudinal instability of relevant networks, particularly those related to mood state. Indeed, such “dense” longitudinal sampling of individual patients has been proposed as an important tool for better understanding the dynamic nature of psychiatric disorders [14].

Several cross-sectional studies have found associations between FC and mood state in individuals with BD. This includes increased mania-related FC between ventral striatum (VS) and midbrain [15], between thalamus and middle frontal gyrus [16], and amongst a network centered around amygdala, midbrain, and frontal cortex [17]. In contrast, depression severity has been associated with less FC between ventral striatum and thalamus [18]. Across studies, FC between ventromedial prefrontal cortex (vmPFC) and amygdala have been correlated with mood state [19]. Taken together, these cross-sectional, group-level findings point to a network, including ventral striatum, midbrain (ventral tegmental area), vmPFC, amygdala, and thalamus, that show mood-related changes in individuals with BD (eFigure 1); interestingly, this network overlaps extensively with regions implicated in reward processing [20]. While findings imply that this network may also show within-subject instability in BD, this hypothesis has not yet been tested.

**Figure 1.**
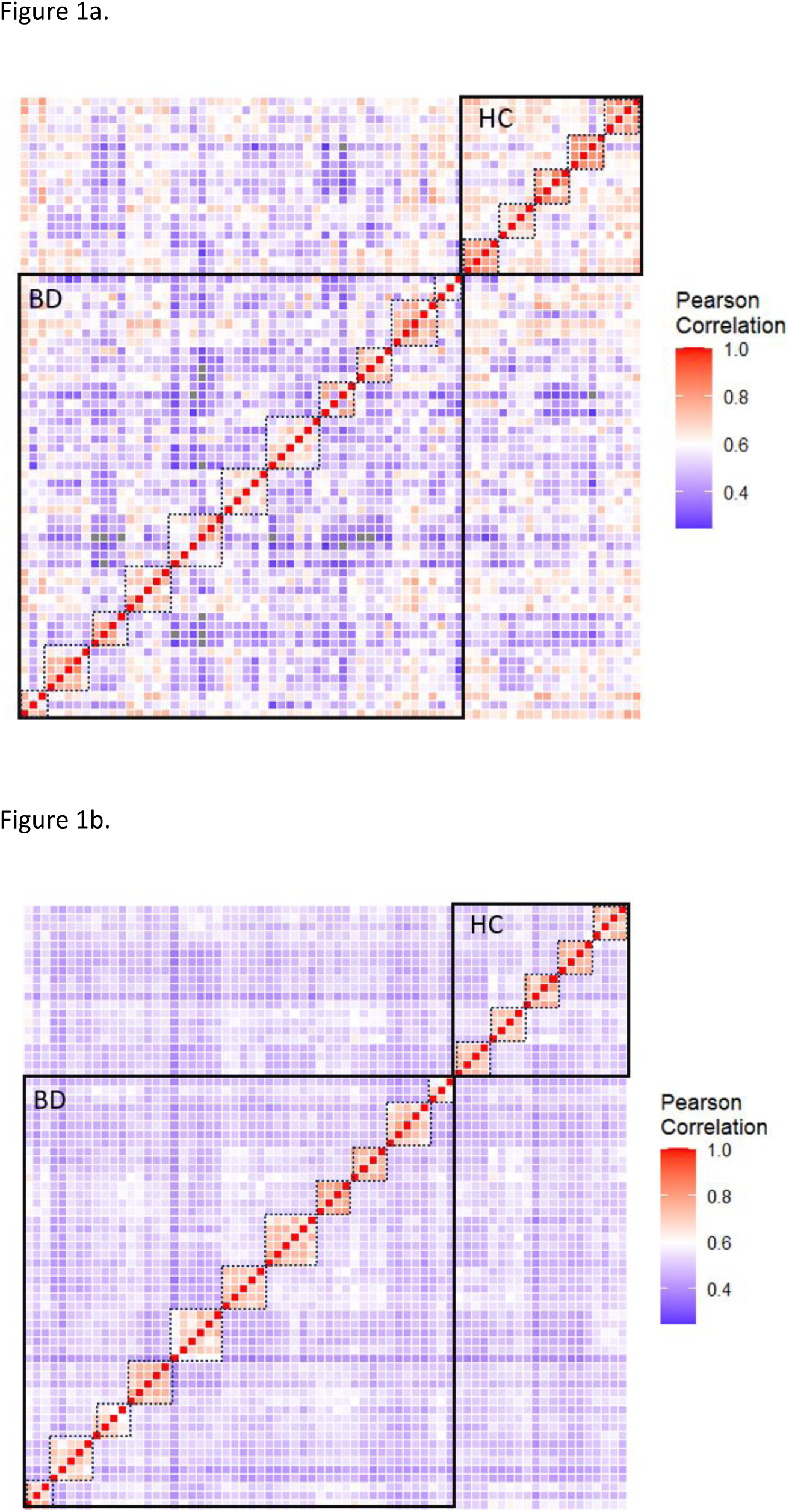
NOI (1a) and Whole Brain (1b) Similarity Across Between- and Within-Subject Scans. Each square represents a scan-pair. Dotted lines demarcate individual subjects. Larger boxes distinguish BD (lower-left) vs. HC (upper-right) participants. Both NOI (1a) and whole-brain (1b) plots show higher within-subject vs. between-subject similarity. Within-subject NOI similarity (1a, within dotted boxes) is lower in BD (lower-left) vs. HC (upper-right).

The current study is the first to utilize precision imaging to assess network stability longitudinally in BD, focusing on circuitry previously found to be associated with mood state (and largely overlapping with a canonical reward-related network). Specifically, we collected 20 minutes of resting-state data in young people with BD 4-6 times over a nine-month window, preferentially scanning during different mood states; HC were scanned four times. We hypothesized that individuals with BD (vs. HC) would show less longitudinal stability in a mood-related network, differences that would be accentuated with hypomania and depression.

## METHODS

### Participants

Young people (age range: 13-30 years old) with BD were recruited from Child and Adolescent Bipolar Services (CABS), an outpatient specialty clinic at the University of Pittsburgh Medical Center. As part of the CABS clinic evaluation, a detailed and comprehensive clinical assessment based on the Kiddie Schedule for Affective Disorders and Schizophrenia – Present/Lifetime (KSADS-PL) [21] was conducted; clinical diagnoses were obtained via an examination of medical records and discussion with the treatment team, and confirmed with the participant at intake. Young people without psychopathology (HC), group-matched on age to the BD participants, were recruited from a research registry at the University of Pittsburgh (Pitt+Me®). The KSADS-PL was used to screen HC for psychopathology and past treatment; family history was also assessed.

To maximize diverse mood states across follow-up, we preferentially recruited currently or recently symptomatic BD participants, specifically those with BD-I/II who had experienced ≥1 depressive episode and ≥1 manic, hypomanic, or mixed episode in the past year. BD participants could not have a primary psychotic disorder or schizophrenia. HC participants could not have a current or past DSM-5 diagnosis, and could not have a first-degree family history of BD. Exclusion criteria for both groups included: pregnant or planning to become pregnant within the next 6 months; history of head injury, neurological or systemic medical disease, or pervasive developmental disorder that could impact MRI scans; substance use disorder within the past year; and the presence of metallic foreign objects.

Informed consent was obtained from all adult participants; for adolescents, informed consent was obtained from a parent/guardian and the adolescent gave assent. All procedures were approved by the University of Pittsburgh Institutional Review Board.

### Procedures (BD)

Following intake, the first MRI scan was scheduled within 1-2 weeks. Within 24 hours prior to the scheduled scan, participants completed an interview that included *(1)* dimensional assessments of manic (Young Mania Rating Scale; YMRS) [22] and depressive symptoms (Montgomery-Åsberg Depression Rating Scale; MADRS) [23]; and *(2)* the Psychiatric Status Rating (PSR) Scale from the Longitudinal Interview Follow-up Evaluation (LIFE) [24] (past three months). Based on PSR depression and hypomania rating lines, mood state over the week prior to scan was classified as *(1)* euthymic (<3 on depression and hypomania); *(2)* depressed (≥3 on depression; <3 on hypomania); *(3)* hypomanic (≥3 on hypomania; <3 on depression); or *(4)* mixed (≥3 on depression and hypomania). Threshold depression and hypomania were defined according to a rating of ≥5 on the depression and hypomania lines, respectively. Medications and substance use over the preceding follow-up period were also recorded. All assessments were conducted by a trained interviewer (JF), who presented to a child psychiatrist for consensus (DMH).

Participants then underwent an hour-long MRI protocol, which included two 5-minute resting-state runs (fixation cross, eyes open), interleaved with two 5-minute Inscapes runs (minimally engaging video that activates similar circuitry to resting state [25]) (total: 20 minutes of resting fMRI). At the scan, participants completed questionnaires to assess state arousal/sleepiness (ABCD Questionnaire [26]) and sleep-related impairment over the past week (PROMIS Sleep-Related Impairment) [27].

Over a 9-month follow-up, participants with BD were scanned 3-5 additional times (using the protocol described above), preferentially during different mood states. To facilitate this, weekly brief self-report questionnaires were administered to assess changes in depressive (Patient Health Questionnaire-9) [28] and manic symptoms (Altman Mania Rating Scale) [29]. Participants were invited for a pre-scan visit if: (1) questionnaires indicated a previously unscanned threshold mood state (PSR ≥5; e.g., hypomania), with ≥2 weeks since previous scan; (2) questionnaires indicated a recurrent scanned threshold mood state (PSR ≥5) or subthreshold episode (PSR=3 or 4) of a previously unscanned polarity (e.g., subthreshold mania), with ≥4 weeks since previous scan; or (3) ≥12 weeks since previous scan (see eFigure 1) . If the above scanning criteria were confirmed based on the PSR depression and hypomania rating lines over the past week, participants were scanned within 24 hours of the scan visit.

### Procedures (HC)

Following confirmation of eligibility at intake, HC participants were scheduled for their initial scan. Scanning procedures were identical to those described above. Following initial scan, an additional 3 scans were scheduled approximately every 12 weeks. HC participants did not complete mood questionnaires; however, they did complete the ABCD and PROMIS questionnaires to assess state arousal/sleepiness and sleep-related impairment, respectively.

### Scanning Parameters

All data were collected on a single Siemens 3T Prisma scanner at the Pitt-CMU Bridge Center using a 64-channel head coil. At the first scan only, high-resolution T1- and T2-weighted images were acquired (T1: MPRAGE, 2.3s TR, 1 x 1 × 1 mm voxels; T2: 3.0s, 1 x 1 x 1 mm voxels). During all scans, functional imaging data were collected using a multiband (MB) gradient-echo EPI sequence (TR = 1.5 s, TE = 30 ms, flip angle = 74, voxel size = 2 x 2 x 2 mm, 68 slices, MB=4). Two acquisitions in opposite field encoding directions were utilized to construct fieldmaps.

### Data Analysis

Details regarding preprocessing steps and parameters can be found in the eMethods. Briefly, initial preprocessing steps were implemented using *fmriprep 20.2.6 LTS*, implemented in Flywheel. Functional images were processed via the following steps: (1) susceptibility distortion correction; (2) co-registration to the T1w reference; (3) slice-time correction; (4) normalization to MNI space; and (5) creation of a confounds file, including motion nuisance regressors and global signal. The following postprocessing steps were conducted using *xcp_d*, implemented within a singularity framework: (1) outlier detection, excluding volumes with low-pass filtered (<.1 Hz) framewise displacement (FD) greater than 0.1mm [30]; (2) despiking, mean-centering, and linear detrending; and (3) nuisance regression using 36 confounds from *fmriprep* output (six motion parameters, global signal, mean white matter signal, and mean CSF signal; temporal derivatives of these parameters; and quadratic expansion of the above). Sessions with >20% data loss due to high motion were removed.

Using nilearn’s NiftiLabelsMasker, we extracted functional timeseries and correlation matrices from residualized BOLD signals for the Shen atlas (268 parcels) [31]. We utilized this atlas because it includes both cortical and subcortical parcels, and has been frequently used in the literature to predict clinical and behavioral outcomes (e.g., [32]). Nilearn scripts were also used to assess coverage; outlier scans with poor coverage were removed. To ensure adequate coverage and minimize variability due to differential within- or between-person coverage of a particular parcel (which can lead to less reliability across scan [33]), we only included parcels with >90% coverage across all participants in final analysis (194/268; 72.4% of parcels included). For the network-of-interest (NOI) analysis, we used Neurosynth [34] (default thresholds) to identify Shen parcels corresponding to the following *a priori* terms: “ventral striatum” [121, 125, 258, 259], “ventral tegmental area” [132, 265], “ventromedial prefrontal cortex” [3, 5, 134, 138], “amygdala” [92, 228], and “thalamus” [126–128, 262–264]. Seventeen of 18 (94%) of parcels had >90% coverage and were included in analysis; parcel 3 was excluded (eFigure 2).

**Figure 2.**
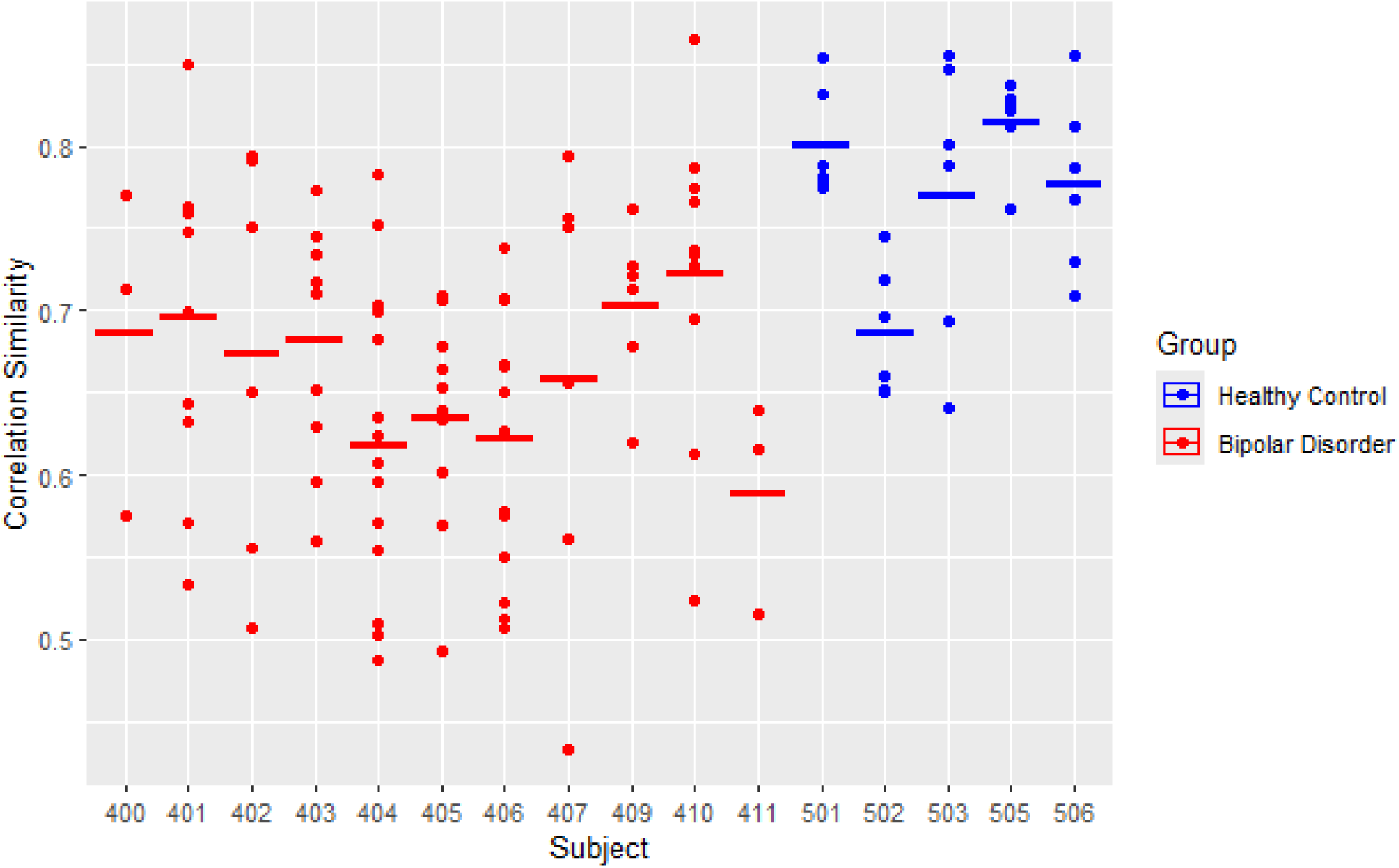
NOI Within-Subject Stability in BD vs. HC Youth. Dots indicate the similarity between each within-subject scan pair; the horizontal line demarcates the median subject-level similarity across scan pairs. HC, shown in blue, have overall higher average NOI stability than BD youth.

In R 4.2.0, we first linearized the upper triangle of the correlation matrix and averaged across resting-state and Inscapes runs (eFigure 3). To test similarity, we assessed within-versus between-person correlation for NOI and whole-brain analyses. Nesting within individual and scan, we then assessed whether BD (vs. HC) within-person scan pairs showed different levels of NOI and whole-brain stability (i.e., Pearson correlation across scans). All scan-pair-level models were adjusted for motion (mean FD), interscan interval, and time-of-day differences. Due to the complex nesting structure, frequentist models did not converge; thus, we used Bayesian models with default priors (R package *rstanarm*) to run models, with group (BD vs. HC) as an independent variable and scan-pair stability as the dependent variable. We report 95% and, where relevant (to assess for marginal associations), 90% prediction intervals (PI). We also assessed person-level relationship between group and average scan-pair stability, using Mann-Whitney U tests due to a limited number of observations. To determine the degree to which scan-pair stability distinguished BD vs. HC, we calculated the Area Under the Receiver Operating Characteristic Curve (AUC) for NOI analysis.

**Figure 3.**
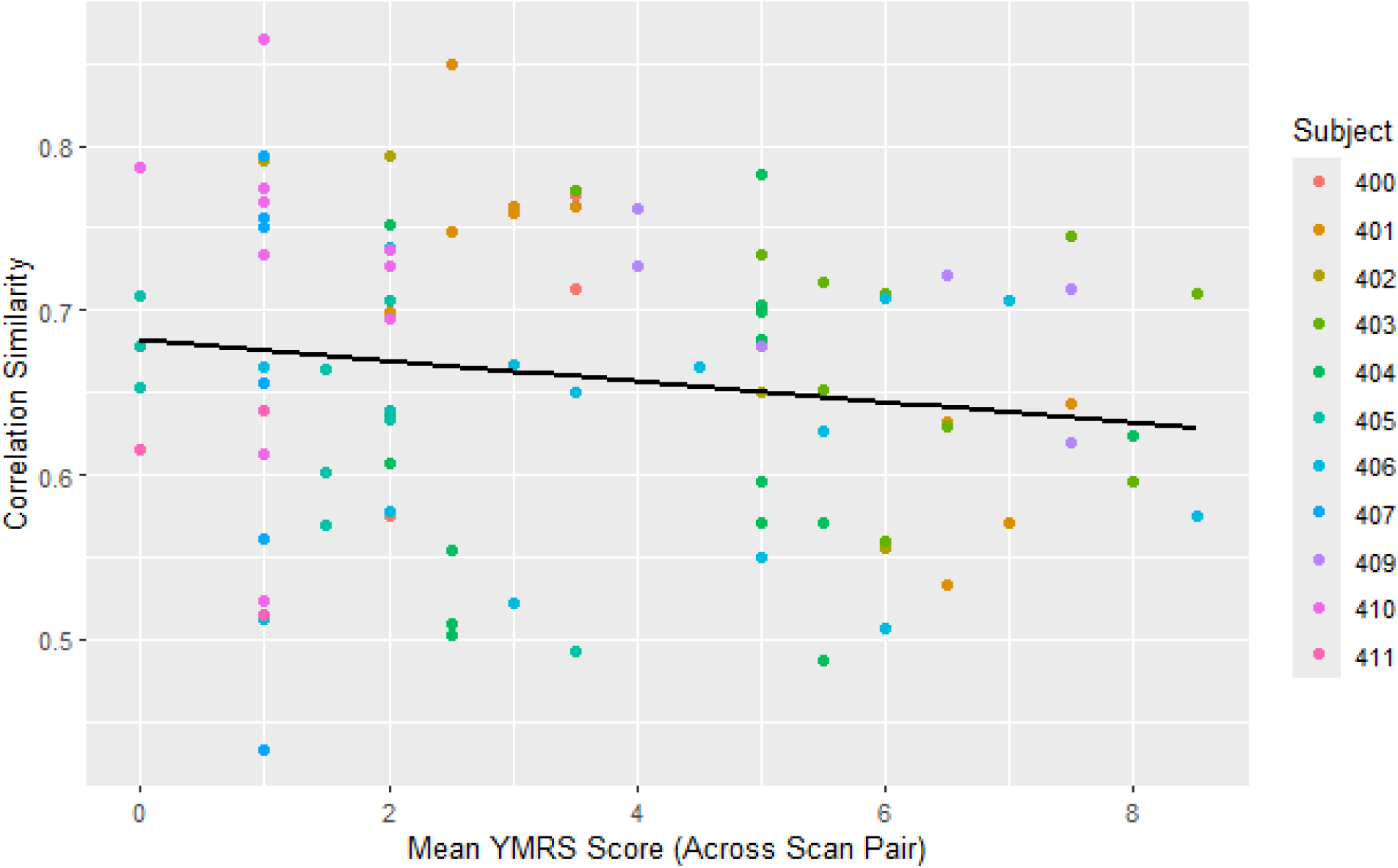
Relationship between scan-pair mean YMRS and NOI stability, grouped by subject.

We also used Bayesian models (as above) to assess the relationship between scan-pair NOI instability and mood symptoms, testing the association between (1) YMRS and MADRS, averaged across scan pair (independent variables) and (2) scan-pair stability (dependent variable). Second, we assessed whether group results (BD vs. HC) persisted after removing scans in relevant mood states (determined by the PSR the week of scan) and when only including mood-congruent scan pairs (i.e. both scans in the same mood state). In addition, Bayesian models were used to assess the effect of BD subtype, testing within the BD sample whether BD subtype (BD-I vs. -II; independent variable) was associated with scan-pair NOI stability (dependent variable).

Further analyses were performed, on the scan-pair and subject level as appropriate, to assess the effect of bipolar subtype, as well as potential confounds including comorbidity, medications, and sleep/arousal. Additionally, we assessed whether findings were specific to the pre-specified NOI as constructed via the Shen atlas, or if they were also found in (1) another subcortical atlas (Tian; [35]) and/or (2) Shen canonical networks. Finally, we assessed whether NOI instability was found *within-scan*, testing (1) split-half and (2) cross-condition (i.e., Rest vs. Inscapes) stability. As above, Bayesian mixed models (nested within subject) were used to assess whether diagnostic group (BD vs. HC; independent variable) was associated with within-scan stability (dependent variable), adjusting for motion, interscan interval, and time-of-day differences.

## RESULTS

### Sample

Twelve young people with BD were recruited, along with six HC. One participant with BD withdrew after a single scan and is not included in analysis; an additional BD scan was excluded due to poor coverage. The remaining eleven participants had a total of 50 high-quality scans: 20 euthymic; 21 depressed (7 threshold); 6 hypomanic (5 threshold); and 3 mixed (all subthreshold) (see Methods for definitions; eFigure 4). Two BD participants only completed three scans due to loss to follow-up. Of the six HC, one participant was excluded due to excessive motion; the remaining 5 participants had a total of 20 high-quality scans. In total, these high-quality scans generated 1,831 between-person scan-pairs and 124 within-person scan-pairs (94 BD, 30 HC). BD vs. HC did not differ according to motion, as measured by mean FD (p=0.6).

Participants were a median of 19.3 years old (range 16.3 to 23.3 years old), 75% female, and majority white (68.8%) (Table 1), with no significant differences in demographic characteristics across groups. Comorbidity in the BD youth was common; 8/11 (72.7%) had an anxiety disorder and 3/11 (27.3%) had ADHD. Most BD participants were on an atypical antipsychotic (10/11; 90.9%), lamotrigine (6/11; 54.5%), and/or an antidepressant (7/11; 63.6%) during at least one scan visit.

**Table 1.** Demographic and Clinical Characteristics of the Sample.

|  | BD (n=11) | HC (n=5) | Total (n=16) |
| --- | --- | --- | --- |
| **Age: Median [Min, Max]** | 19.2 [17.1, 23.3] | 19.4 [16.3, 21.2] | 19.3 [16.3, 23.3] |
| **Sex: n (%) Female** | 8 (72.7%) | 4 (80.0%) | 12 (75.0%) |
| **Gender Identity n (%) Women** | 6 (54.5%) | 4 (80.0%) | 10 (62.5%) |
| **Race** |  |  |  |
| *Asian* | 2 (18.2%) | 1 (20.0%) | 3 (18.8%) |
| *White* | 7 (63.6%) | 4 (80.0%) | 11 (68.8%) |
| *Biracial* | 2 (18.2%) | 0 (0%) | 2 (12.5%) |
| **Ethnicity** |  |  |  |
| *Hispanic* | 1 (9.1%) | 0 (0%) | 1 (6.3%) |
| *Non-Hispanic* | 10 (90.9%) | 5 (100%) | 15 (93.8%) |
| **Bipolar Disorder Subtype** |  |  |  |
| *Bipolar I* | 6 (54.5%) |  |  |
| *Bipolar II* | 5 (45.5%) |  |  |
| **Medications** |  |  |  |
| *Lithium* | 4 (36.4%) |  |  |
| *Lamotrigine* | 5 (45.5%) |  |  |
| *Atypical Antipsychotic* | 10 (90.9%) |  |  |
| *Antidepressant* | 7 (63.6%) |  |  |
| *Benzodiazepine* | 4 (36.4%) |  |  |
| *Stimulant* | 3 (27.3%) |  |  |
| **Comorbid Disorders** |  |  |  |
| *ADHD* | 3 (27.3%) |  |  |
| *GAD* | 6 (54.5%) |  |  |
| *Panic Disorder* | 1 (9.1%) |  |  |
| *Social Anxiety Disorder* | 4 (36.4%) |  |  |
| *PTSD* | 2 (18.2%) |  |  |
| *OCD* | 2 (18.2%) |  |  |

BD Bipolar Disorder; HC Healthy Control; ADHD Attention-Deficit Hyperactivity Disorder; GAD Generalized Anxiety Disorder; PTSD Post-Traumatic Stress Disorder; OCD Obsessive- Compulsive Disorder

### Within-vs. Between-Person Scan-Pair Similarity (Fig 1)

Across the entire sample, average NOI across-scan similarity was higher within-person (r=0.70) than between-person (r=0.54) (t=8.05, p<.0001). Replicating previous work [10], average whole-brain across-scan connectome similarity was higher within-person (r=0.71) than between-person (r=0.51) (t=19.9, p<.0001).

### Within-Person Scan-Pair Stability in BD vs. HC

Within the mood-related circuitry NOI, BD youth (vs. HC) showed lower scan-pair stability (mean -0.109; 95% PI -0.181, -0.038) (Figs 1a, 2). This strength of this relationship was not significantly impacted by sex or age (eResults: Demographic Characteristics). On average, BD youth (vs. HC) showed lower scan-pair stability (Mann-Whitney U p=.003) and NOI scan-pair stability distinguished youth with BD vs. HC with excellent accuracy (AUC=0.95). The optimal threshold for distinguishing BD vs. HC (based on Youden’s index) was 0.75 (specificity=0.8, sensitivity=1). There were no substantial between-group (BD vs. HC) differences in whole-brain scan-pair stability (mean -0.031; 95% PI -0.08, 0.014; Mann-Whitney U p=.18) (Fig 1b).

### Mood Symptoms/State and Within-Person Scan-Pair Stability

Within BD youth, more manic symptoms (i.e., average of YMRS across scan pair) was associated with lower NOI scan-pair stability (mean -0.012; 95% PI -0.023, -0.0002) (Fig 3). There was no appreciable relationship between depressive symptoms across scans (measured via MADRS) and NOI stability (mean .002; 90% PI -0.001, 0.006). Whole-brain scan-pair stability was not associated with depressive or manic symptoms within BD youth (YMRS: mean -0.002; 90% PI -0.009, 0.004; MADRS: mean 0.000; 90% PI -0.002, 0.002).

After removing scans classified as hypomanic or mixed (35/94 BD scan pairs removed), BD (vs. HC) still showed less NOI scan-pair stability, though the effect size was slightly attenuated (mean -0.089; 95% PI -0.170, -0.006). On the subject level, after excluding hypomanic or mixed scans, BD youth (vs. HC) also showed significantly less within-person stability (Mann-Whitney U p=.02); and still distinguished BD vs. HC youth with very good accuracy (AUC=.87). These findings also persisted after removing scan pairs with discordant mood states (60/94 BD scan pairs) (mean -0.1; 95% PI -0.183, -0.017), indicating that NOI instability was not attributable to across-scan mood changes.

### Effect of Bipolar Subtype

To assess the effect of BD subtype, we first assessed whether BD-I differed from BD-II. We found that BD-II showed marginally less NOI scan-pair stability than BD-I (mean -0.071; 95% PI -0.149, 0.006; 90% PI -0.136, -0.007). Compared to HC, both BD-I and BD-II groups showed less scan-pair stability, though the effect was larger in BD-II (BD-I vs. HC: mean -0.078; 95% PI -0.148, -0.006; BD-II vs. HC: mean -0.148; 95% PI -0.222, -0.075) (eFigure 5).

### Sensitivity Analyses: Potential Confounds

#### Comorbid Disorders

At the subject level, we assessed whether group differences in NOI stability persisted after sequentially removing youth with each comorbidity listed in Table 1. All models remained significant (Mann-Whitney U p-values 0.003 to 0.02), indicating that group-level differences (BD vs. HC) were not driven by a comorbid disorder. At the scan-pair level, we assessed whether comorbid diagnoses had an independent effect on NOI stability across the sample, including BD status in the model. All 90% PIs (for each comorbid disorder, as well as for combined anxiety disorders) included the null, while the 95% PIs for BD in each model excluded the null (eTable 1). While substance use disorder in the past year was exclusionary, four participants had subthreshold substance use (cannabis or alcohol) over at least one follow-up period (5 scans). After removing scans following a period of substance use, findings were still significant (mean -0.096; 95% PI -0.168, -0.02).

#### Medications

To test the effect of medications, we assessed both across-scan medication changes and individual medication classes at time of scan (listed in Table 1). Within the BD sample, we did not observe an association of NOI stability with across-scan medication changes (mean -0.006; 90% PI -0.024, 0.012). After removing BD scan-pairs with discordant medications (51/94 BD scan-pairs excluded), we still found that BD (vs. HC) showed lower NOI stability (mean -0.096; 95% PI -0.177, -0.006), indicating that results were not explained by across-scan medication changes. To test the effect of medication classes on NOI stability, we included exposure to medication class (in one or both scans) as a covariate and, in separate models, assessed the effects of each medication class. Across models, all 90% PIs for each medication class included the null; and BD remained associated with decreased NOI stability (95% PIs did not include the null) (eTable 2).

#### Sleep and Arousal

To test effects of changes in sleep and arousal, we assessed *(1)* observed sleepiness during scan (defined as >5 seconds of eyes closed, assessed via eye-tracking), *(2)* state sleepiness at the time of scan (average of “tiredness” and “sleepiness” items on the ABCD questionnaire), and *(3)* sleep-related impairment over the past week (PROMIS).

Sleepiness was observed during at least one scan of 15/94 (16.0%) of BD scan-pairs and 9/30 (30%) of HC scan-pairs. Including sleepiness (yes/no) in the model, BD remained a significant predictor of lower NOI stability (mean -0.115; 95% PI -0.181, -0.047); sleepiness was also marginally associated with lower NOI stability (mean -0.054; 90% PI -0.102, -0.008). In whole-brain analysis, observed sleepiness was also associated with lower stability (mean - 0.042; 95% PI -0.078, -0.006). Regarding self-report measures of sleep-related impairment and arousal at scan, there was no relationship between these measures on NOI stability (90% PIs included the null) and, even after adjustment for these variables, the effect of BD remained (95% PIs excluding the null).

#### Methodological Factors

We conducted additional sensitivity analyses to ensure that observed findings were not due to (1) variable number of scan pairs in the BD youth or (2) the exclusion of Shen parcels with incomplete (<90%) coverage. Findings persisted even when these factors were accounted for (eResults).

### Supplemental Analyses

#### Specificity of Findings to the Shen Atlas

To assess specificity of findings to the selected NOI, we conducted stability analyses in anatomically analogous nodes of the subcortical Tian parcellation (Tian, scale 1; [35]). Utilizing the Tian parcellation, BD (vs. HC) still showed lower NOI stability (mean -0.08; 95% PI -0.16, -0.005), indicating that findings were not parcellation-specific (eResults).

#### Specificity of Findings to the Hypothesized NOI

To assess specificity of findings to the selected NOI, we conducted stability analyses in canonical Shen networks. BD vs. HC showed lower stability in the Cerebellum (mean -0.12; 95% PI -0.235, -0.006) and, to a lesser extent, in the Basal Ganglia (mean -0.093; 90% PI -0.166, -0.015) networks (eTable 3). However, these findings did not distinguish BD vs. HC youth as accurately as the hypothesized NOI (AUCs=0.76, 0.80; Mann-Whitney p-values>.05) (eResults). Thus, instability may not be specific to the hypothesized NOI, but also found to a lesser degree in a subset of other (subcortical) networks.

#### Stability Within Scan

To test whether NOI instability was also present within-scan, we assessed split-half and cross-condition (i.e., Rest vs. Inscapes) stability. We found that both were significantly lower in BD vs. HC (split-half NOI stability: mean -0.073; 95% PI -0.129, -0.018; cross-condition NOI stability: mean -0.078; 95% PI -0.147, -0.01). Because previous work has found within-scan stability to relate to arousal, we adjusted for observed sleepiness; this did not impact findings (eResults). BD (vs. HC) still showed lower scan-pair instability in the mood-related NOI, even after adjusting for within-scan stability measures, indicating that within-scan stability did not explain the primary observed relationship (eResults). Whole-brain, within-scan stability did not differ across groups.

## DISCUSSION

In a sample of 70 scans, collected from 11 youth with BD and 5 HCs, we assessed NOI stability as a possible neural correlate of BD. We found that, similar to previous work, both whole-brain and NOI FC were relatively stable within-person (average correlation ∼0.7). FC within a mood-related NOI (and to some extent subcortical networks, but not the whole brain) was less stable in BD youth vs. HCs. FC NOI in particular distinguished the 11 BD youth (especially youth with BD-II) from 5 HC. Manic symptoms, but not depressive symptoms, were marginally correlated with less NOI stability, although diagnostic group differences persisted even after removing predominantly manic/hypomanic scans. Results did not appear to be explained by potential confounds, including time-of-day differences, time between scans, motion, comorbidities, medications, sleep and arousal, or methodological factors; however, scan-pairs with observed sleepiness during at least one scan showed less stability over time. Interestingly, BD youth (vs. HC) also showed lower within-scan (split-half and cross-condition) stability in the NOI, indicating that instability may be observable on a range of timescales; however, within-scan stability did not explain the observed relationship between BD and NOI scan-pair instability.

For network instability to be a potential person-level marker of BD, an important prerequisite is that networks are longitudinally stable in HC. While test-retest reliability of individual edges obtained from resting-state data using standard acquisition times (e.g., 5-minutes) remains fairly low [36], the growing field of precision imaging has demonstrated that within-person stability is enhanced with increased data acquisition times and advanced denoising strategies [10]. Consistent with this work, we find that whole-brain networks are stable across time in both BD and HC (r∼.7), and within vs. between-subject scan pairs show higher correlations. Our findings add to this work by demonstrating that even a smaller, subcortical network (mood-related NOI) shows similar within-subject stability (r∼.7), which is similarly higher than between-subject correlation, thus presenting an opportunity to assess NOI instability as a potential person-level neural marker of BD. This stability is in contrast to reward task activation, which shows lower test-retest reliability (i.e. ICCs<0.5), particularly in a reward network (that overlaps substantially with the mood-related network assessed here) [37]; thus, while a reward task reliably activates the network examined here, this activation does not demonstrate the necessary longitudinal stability in healthy populations to directly assess within-person instability in BD.

The current approach also overlaps with the field of functional connectome fingerprinting, which indicates a high within-subject (vs. between-subject) similarity across resting-state scans, allowing for the identification of a subject’s second scan based on their first, even in large samples [38]. Functional connectome stability may increase across development, and delays in this increase, both brain-wide and within specific networks, have been associated with psychopathology [13] and lower social skills [12]. In addition, a recent study found that lower fingerprinting accuracy within the cingulo-opercular network over a 4-month period also predicted psychological distress in adolescents [39]. This work points to longitudinal instability, particularly in networks of interest, as a possible and novel marker of psychopathology. To our knowledge, the current study is the first to test and demonstrate such longitudinal instability in BD.

We hypothesized that a mood-related NOI would show longitudinal within-person instability in BD youth due to previously observed mood-related correlations in cross-sectional studies. Consistent with our hypotheses, we found an effect of mood state, such that scan-pairs with a higher level of manic symptoms showed a greater extent of NOI instability. However, BD vs. HC still showed less NOI stability even when scans with manic symptoms were removed, indicating that this may also be a trait marker of BD. These mood-state independent (i.e., trait) findings highlight the possibility that NOI instability may not just be a consequence of mood change but may also be related to the *propensity* for hypomania (even during euthymic mood). This is consistent with the computational theory that BD is related to an underlying oscillation of networks [40] coupled with positive feedback loops [41] that give rise to mood episodes. Interestingly, NOI instability was not related to depressive symptoms, indicating that this marker may underlie manic symptoms more specifically. Such specificity also provides some indication that this marker may be particular to BD (as opposed to found in unipolar depression as well); however, this hypothesis should be tested in future studies.

Interestingly, we found that, while both BD-I and -II showed less NOI stability than HC, the effect was strongest in BD-II. While this may seem counterintuitive if we consider BD-I to be more severe than BD-II, it is consistent with work that shows that, in tertiary referral centers, BD-II patients are in some ways more severely affected than BD-I patients; for example, they are more likely to have rapid cycling, ≥10 prior mood episodes, and worse overall functioning [42]. The finding is also in line with earlier work indicating that BD-II patients are more likely to have mood lability and neuroticism, compared to both BD-I and unipolar depressed [43]. While extremely preliminary, the possibility that NOI instability may be especially prominent in BD-II highlights potential clinical utility, given that the differential diagnosis of BD-II (vs. unipolar depression) can be quite challenging.

We conducted several sensitivity analyses to assess the effects of potential confounds on network instability, including medications, comorbid disorders, and sleep/arousal. Of the tested covariates, the only predictor of network instability (both NOI and whole-brain) was observed sleepiness during scan. This network instability is consistent with previous work that has found within-scanner sleep to be associated with lower within-DMN connectivity and altered thalamo-cortical connectivity [44]. In the current study, observed sleepiness was not associated with diagnostic group or mood symptoms, and adjustment for this covariate did not impact results; in fact, the mean group effect was slightly increased with adjustment. However, future studies of network instability should take this potential confound into account. Of note, self-report measures of arousal and sleepiness do not show this relationship to network instability, so the careful documentation of closed eyes appears to be critical.

Results should be interpreted in the context of the following limitations. First, although our analyses included 70 high-quality 20-minute fMRI scans, we only included 11 BD youth and 5 HC. While recent high-impact precision imaging studies have been published with even fewer participants [45, 46], such an approach limits the assessment of person-level confounds and potentially the generalizability of results. We also had a small number of scans during hypomanic or mixed states, limiting the assessment of the impact of hypomanic symptoms. In addition, in the setting of an underpowered pilot study, interpretation of negative findings is limited. Thus, a future study with a larger number of participants will be necessary to rigorously assess this marker in a racially, ethnically, and clinically diverse sample. Second, as a first pass, we compared BD to HC. A critical next step would be to assess whether it specifically distinguishes BD from other disorders (e.g., unipolar depression). Third, we selected participants with at least one episode of each polarity in the past year and scanned them preferentially during different mood states. While such an approach was critical to assess the relationship between NOI stability and mood symptoms, this finding may not hold in a sample of relatively asymptomatic participants with BD. Fourth, most of our participants were female and, given recent work demonstrating that FC may fluctuate with the menstrual cycle [47], this is an important confound will be measured in the future; however, given the lack of an observed relationship between sex and NOI stability, it is unlikely that this factor would explain observed group (BD vs. HC) differences. Fifth, while we did not observe an effect of medication, we did not collect information about as-needed medications or medication adherence; thus, it is possible that these factors may have confounded observed results.

In conclusion, this study is (to our knowledge) the first to apply precision imaging methods to BD, scanning youth with BD multiple times (preferentially in mood episodes) over a 9-month period, along with HC. Results indicate that instability in a mood-related network may be a potential neural marker for BD, with a person-level interpretation. While additional studies are necessary to assess the specificity and generalizability of this measure, this novel approach may shed light on the neural origins of mood instability in BD.

## Supporting information

eSupplement

## AUTHOR CONTRIBUTIONS

Dr. Hafeman made substantial contributions to the conception and design of this work; the acquisition, analysis, and interpretation of the data; and drafting and revising the manuscript critically for intellectual content. Ms. Feldman and Ms. Mak made substantial contributions to the acquisition of the data; and revising the work critically. Mr. Merranko and Dr. Gratton made significant contributions to the analysis and interpretation of the data; and revising the work critically. Drs. Goldstein, Phillips, and Birmaher made substantial contributions to the interpretation of the data; and revising the work critically for intellectual content. All authors approve the final version to be published and agree to be accountable for all aspects of the work, ensuring that questions regarding accuracy or integrity have been addressed.

## ACKNOWLEDGEMENTS

The authors would like to acknowledge the study participants. This work was supported by the Brain and Behavior Research Foundation (2019 NARSAD Young Investigator Award; PI: Hafeman), the Galena-Yorktown Foundation, and K23MH110421 (PI: Hafeman). This research was supported in part by the University of Pittsburgh Center for Research Computing, RRID:SCR_022735, through the resources provided. Specifically, this work used the HTC cluster, which is supported by NIH award number S10OD028483.

