## Supplementary material for "Longitudinal Stability of Mood-Related Resting-State Networks in Youth with Symptomatic Bipolar-I/II Disorder": eSupplement

**eSupplement: Table of Contents**

*eMethods: p. 1*

*eResults: p. 5*

*eTables: p. 8*

*eFigures: p. 10*

*References: p. 14*

**eMethods:**

Preprocessing and Data Analysis: The T1-weighted (T1w) image was corrected for intensity non-uniformity (INU) with N4BiasFieldCorrection (Tustison et al. 2010), distributed with ANTs 2.3.3 (Avants et al. 2008, RRID:SCR_004757), and used as T1w-reference throughout the workflow. The T1w-reference was then skull-stripped with a Nipype implementation of the antsBrainExtraction.sh workflow (from ANTs), using OASIS30ANTs as target template. Brain tissue segmentation of cerebrospinal fluid (CSF), white-matter (WM) and gray-matter (GM) was performed on the brain-extracted T1w using fast (FSL 5.0.9, RRID:SCR_002823, Zhang, Brady, and Smith 2001). Brain surfaces were reconstructed using recon-all (FreeSurfer 6.0.1, RRID:SCR_001847, Dale, Fischl, and Sereno 1999), and the brain mask estimated previously was refined with a custom variation of the method to reconcile ANTs-derived and FreeSurfer-derived segmentations of the cortical gray-matter of Mindboggle (RRID:SCR_002438, Klein et al. 2017). Volume-based spatial normalization to two standard spaces (MNI152NLin2009cAsym, MNI152NLin6Asym) was performed through nonlinear registration with antsRegistration (ANTs 2.3.3), using brain-extracted versions of both T1w reference and the T1w template. The following templates were selected for spatial normalization: ICBM 152 Nonlinear Asymmetrical template version 2009c [Fonov et al. (2009), RRID:SCR_008796; TemplateFlow ID: MNI152NLin2009cAsym] and FSL’s MNI ICBM 152 non-linear 6th Generation Asymmetric Average Brain Stereotaxic Registration Model [Evans et al. (2012), RRID:SCR_002823; TemplateFlow ID: MNI152NLin6Asym].

Functional data preprocessing

For each of the BOLD runs found per subject (across all tasks and sessions), the following preprocessing was performed. First, a reference volume and its skull-stripped version were generated using a custom methodology of fMRIPrep. A B0-nonuniformity map (or fieldmap) was estimated based on two (or more) echo-planar imaging (EPI) references with opposing phase-encoding directions, with 3dQwarp Cox and Hyde (1997) (AFNI 20160207). Based on the estimated susceptibility distortion, a corrected EPI (echo-planar imaging) reference was calculated for a more accurate co-registration with the anatomical reference. The BOLD reference was then co-registered to the T1w reference using bbregister (FreeSurfer) which implements boundary-based registration (Greve and Fischl 2009). Co-registration was configured with six degrees of freedom. Head-motion parameters with respect to the BOLD reference (transformation matrices, and six corresponding rotation and translation parameters) are estimated before any spatiotemporal filtering using mcflirt (FSL 5.0.9, Jenkinson et al. 2002). BOLD runs were slice-time corrected to 0.696s (0.5 of slice acquisition range 0s-1.39s) using 3dTshift from AFNI 20160207 (Cox and Hyde 1997, RRID:SCR_005927). The BOLD time-series (including slice-timing correction when applied) were resampled onto their original, native space by applying a single, composite transform to correct for head-motion and susceptibility distortions. These resampled BOLD time-series will be referred to as preprocessed BOLD in original space, or just preprocessed BOLD. The BOLD time-series were resampled into several standard spaces, correspondingly generating the following spatially-normalized, preprocessed BOLD runs: MNI152NLin2009cAsym, MNI152NLin6Asym. First, a reference volume and its skull-stripped version were generated using a custom methodology of fMRIPrep. Several confounding time-series were calculated based on the preprocessed BOLD: framewise displacement (FD), DVARS and three region-wise global signals. FD was computed using two formulations following Power (absolute sum of relative motions, Power et al. (2014)) and Jenkinson (relative root mean square displacement between affines, Jenkinson et al. (2002)). FD and DVARS are calculated for each functional run, both using their implementations in Nipype (following the definitions by Power et al. 2014). The three global signals are extracted within the CSF, the WM, and the whole-brain masks. The head-motion estimates calculated in the correction step were also placed within the corresponding confounds file. The confound time series derived from head motion estimates and global signals were expanded with the inclusion of temporal derivatives and quadratic terms for each (Satterthwaite et al. 2013). Frames that exceeded a threshold of 0.5 mm FD or 1.5 standardised DVARS were annotated as motion outliers.

All resamplings can be performed with a single interpolation step by composing all the pertinent transformations (i.e. head-motion transform matrices, susceptibility distortion correction when available, and co-registrations to anatomical and output spaces). Gridded (volumetric) resamplings were performed using antsApplyTransforms (ANTs), configured with Lanczos interpolation to minimize the smoothing effects of other kernels (Lanczos 1964). Non-gridded (surface) resamplings were performed using mri_vol2surf (FreeSurfer).

Post-processing of fmriprep outputs

The eXtensible Connectivity Pipeline (XCP) (Ciric et al. 2018; Satterthwaite et al. 2013) was used to post-process the outputs of fmriprep version 20.2.6 (Esteban et al. 2019, 2020, RRID:SCR_016216). XCP was built with Nipype 1.8.5 (Gorgolewski et al. 2011, RRID:SCR_002502). For each of the BOLD series found per subject (across all tasks and sessions), the following post-processing was performed. First, outlier detection was performed. In order to identify high-motion outlier volumes, the six translation and rotation head motion traces were low-pass filtered below 6.0 breaths-per-minute, based on Fair et al. (2020) and Gratton et al. (2020). Next, framewise displacement was calculated using the formula from Power et al. (2014), with a head radius of 50 mm. Volumes with filtered framewise displacement greater than 0.1 mm were flagged as outliers and excluded from nuisance regression (Power et al. 2014). The filtered versions of the motion traces and framewise displacement were not used for denoising. Before nuisance regression, but after censoring, the BOLD data were despiked, mean-centered, and linearly detrended. In total, 36 nuisance regressors were selected from the nuisance confound matrices of fMRIPrep output. These nuisance regressors included six motion parameters, global signal, the mean white matter, the mean CSF signal with their temporal derivatives, and the quadratic expansion of six motion parameters, tissues signals and their temporal derivatives (Ciric et al. 2017; Satterthwaite et al. 2013). These nuisance regressors were regressed from the BOLD data using linear regression - as implemented in Scikit-Learn 0.24.2 (Pedregosa et al. 2011). Any volumes censored earlier in the workflow were then interpolated in the residual time series produced by the regression. The interpolated timeseries were then band-pass filtered to retain signals within the 0.009-0.08 Hz frequency band. The processed BOLD was smoothed using Nilearn with a gaussian kernel size of 8.0 mm (FWHM).

Many internal operations of XCP use TemplateFlow version 0.6.3 (Ciric et al. 2022), Nibabel version 4.0.2 (Brett et al. 2022), numpy version 1.18.5 (Harris et al. 2020), and scipy version 1.9.3 (Virtanen et al. 2020). For more details, see the xcp_d website <https://xcp-d.readthedocs.io>.

*Copyright Waiver: The above boilerplate text was automatically generated by fMRIPrep and XCP with the express intention that users should copy and paste this text into their manuscripts unchanged. It is released under the CC0 license*.

**eResults:**

The following sensitivity analyses were performed to evaluate the impact of potential methodological confounds.

*Demographic Characteristics:* We assessed the degree to which age and sex may have impacted the observed association. First, we ran a model adjusting for age and sex, in addition to the session-pair covariates; the relationship between BD and scan-pair instability was not substantially changed (mean -0.113; 95% PI -0.189, -0.039) and neither age nor sex were associated with scan-pair instability (age: mean 0.007; 90% PI -0.012, 0.026, sex: mean 0; 90% PI -0.065, 0.067). Next, we assessed for interactions between BD and demographics, to assess whether the relationship between BD and NOI scan-pair stability may differ according to sex or age. Interaction terms did not approach significance (BD*age: mean -0.013; 90% PI -0.038, 0.01, BD*sex: mean 0.042; 90% PI -0.089, 0.179).

*Variable number of scans:* While all HC had four scans, BD youth had 3-6 scans. To ensure that this inconsistency was not driving observed results, we assessed scan-pair stability using only the first four scans for each BD participant, removing BD participants (n=2) who had <4 scans. Within this subset (36 BD scans, 20 HC scans), BD (vs. HC) still showed less NOI scan-pair stability (mean -0.083; 95% PI -0.16, -0.0005); and NOI scan-pair stability distinguished groups (9 BD, 5 HC) with excellent accuracy (AUC=.93).

*Exclusion of parcels with coverage <90%:* To improve across-scan reliability, we only included parcels with >90% coverage in all participants. To ensure that this decision did not explain observed results, we reran analyses including all parcels, regardless of coverage. For the NOI, this included right ventromedial prefrontal cortex (Shen parcel 3; 69% coverage). Including this parcel, BD (vs. HC) was still associated with lower NOI stability (mean -0.095; 95% PI -0.165, -0.03), distinguishing groups with good accuracy (AUC=.96).

*Specificity of Findings to the Shen Atlas:* To determine whether findings were specific to the Shen atlas, we tested whether NOI instability was also observed using the Tian subcortical atlas (scale 1). We included all parcels within this atlas that showed substantial overlap (≥25%) with the mood-related mask generated in Neurosynth (10 out of 16 parcels). As described in the Methods, Bayesian models were used to assess the relationship between NOI stability (dependent variable) and BD (independent variable), adjusting for motion (mean FD), time between scans (days), and time of day differences. Within this NOI, BD (vs. HC) showed lower stability (mean -0.082; 95% PI -0.163, -0.005).

*Specificity of Findings to the Hypothesized NOI:* We first tested the overlap (Jaccard similarity) between each canonical Shen network and our primary NOI. Next, we sequentially ran Bayesian models (analogous to the primary analysis) with BD as the independent variable and Shen network scan-pair instability as the dependent variable. Findings are described in eTable 3 and shown in eFigure 6. Across-scan stability of the Cerebellar and, to a lesser extent, Basal Ganglia networks was lower in BD vs. HC (Cerebellar: mean -0.12; 95% PI -0.235, -0.006. Basal Ganglia: mean -0.093; 90% PI -0.166, -0.015). There were no differences between BD vs. HC in other canonical Shen networks. Analogous to the primary NOI analysis, we assessed whether BD (vs. HC) were distinguished by Cerebellar and Basal Ganglia network stability. We found that average stability in these networks showed only trend-level differences between BD and HC, and did not reliably distinguish these groups (Cerebellum: Mann-Whitney p=0.11, AUC=0.76; Basal Ganglia: Mann-Whitney p=0.07, AUC=0.8).

Of note, both the Cerebellum and Basal Ganglia Networks are subcortical brain regions. While there is some evidence that these networks may also be important to mood switches in BD (Claeys et al., 2022), these networks also have lower signal-to-noise ratio and lower within-person stability (eFigure 6; Marek et al., 2018; Greene et al., 2020). Thus, we interpret these findings with caution pending larger studies.

*Within-Scan Stability:* We tested two measures of within-scan stability: (1) split-half stability and (2) cross-condition stability. Runs were conducted in the following order: inscapes1, rest1, movie (8 minutes, not included in this analysis), rest2, and inscapes2. Split-half stability was defined as the correlation between the first (inscapes1, rest1) vs second half of the scan (inscapes2, rest2); cross-condition stability was calculated as the correlation between Inscapes and Rest. Split-half and cross-condition stability measures were highly correlated with each other, even after accounting for BD group (partial r=0.76, p<.001). In contrast, after accounting for BD group, neither of the within-scan stability measures were associated with scan-pair stability (split-half: partial r=0.20, p=.48; cross-condition: partial r=0.37, p=.18). The relationship between NOI scan-pair stability and BD persisted even after adjustment for within-scan stability measures (mean -0.1; 95% PI -0.176, -0.027).

Because of previous work indicating that within-scan stability may be related to arousal, we tested whether findings persisted after adjusting for observed sleepiness during scan. NOI within-scan stability still differed across groups (mean -0.07; 95% PI -0.124, -0.017) and arousal was not associated with NOI within-scan stability (mean -0.001; 90% PI -0.006, 0.005).

**eTable 1:** Effects of adjustment for each comorbidity, on the session level. Prediction Intervals excluding the null are in bold.

| Model | Predictors | Mean | 95% Prediction Interval | 90% Prediction Interval |
| --- | --- | --- | --- | --- |
| 1 | *Bipolar Disorder* | **-0.111** | **-0.189, -0.031** | **-0.177, -0.047** |
|  | *ADHD* | 0.004 | -0.1, 0.105 | -0.082, 0.089 |
| 2 | *Bipolar Disorder* | **-0.113** | **-0.2, -0.031** | **-0.183, -0.047** |
|  | *GAD* | 0.006 | -0.069, 0.087 | -0.056, 0.069 |
| 3 | *Bipolar Disorder* | **-0.11** | **-0.185, -0.034** | **-0.172, -0.047** |
|  | *Social Anxiety* | 0.003 | -0.082, 0.084 | -0.067, 0.071 |
| 4 | *Bipolar Disorder* | **-0.114** | **-0.186, -0.042** | **-0.174, -0.054** |
|  | *Panic* | 0.034 | -0.096, 0.165 | -0.071, 0.137 |
| 5 | *Bipolar Disorder* | **-0.116** | **-0.193, -0.041** | **-0.18, -0.055** |
|  | *PTSD* | 0.027 | -0.072, 0.127 | -0.054, 0.108 |
| 6 | *Bipolar Disorder* | **-0.102** | **-0.174, -0.031** | **-0.162, -0.043** |
|  | *OCD* | -0.041 | -0.132, 0.054 | -0.116, 0.039 |
| 7 | *Bipolar Disorder* | **-0.099** | **-0.193, -0.003** | **-0.177, -0.02** |
|  | *Anxiety** | -0.014 | -0.099, 0.07 | -0.086, 0.057 |

*GAD, Social Anxiety, Panic, and/or PTSD

**eTable 2.** Effects of adjustment for each medication class, on the session level. Prediction Intervals excluding the null are in bold.

| Model | Predictors | Mean | 95% Prediction Interval | 90% Prediction Interval |
| --- | --- | --- | --- | --- |
| 1 | *Bipolar Disorder* | **-0.119** | **-0.214, -0.018** | **-0.199, -0.034** |
|  | *Antipsychotic* | 0.011 | -0.064, 0.088 | -0.053, 0.075 |
| 2 | *Bipolar Disorder* | **-0.104** | **-0.18, -0.025** | **-0.168, -0.039** |
|  | *Lithium* | -0.015 | -0.095, 0.061 | -0.083, 0.047 |
| 3 | *Bipolar Disorder* | **-0.108** | **-0.186, -0.033** | **-0.169, -0.047** |
|  | *Lamotrigine* | -0.006 | -0.066, 0.057 | -0.057, 0.046 |
| 4 | *Bipolar Disorder* | **-0.103** | **-0.19, -0.017** | **-0.172, -0.033** |
|  | *Antidepressant* | -0.012 | -0.083, 0.059 | -0.072, 0.044 |
| 5 | *Bipolar Disorder* | **-0.101** | **-0.175, -0.028** | **-0.162, -0.041** |
|  | *Stimulant* | -0.05 | -0.119, 0.02 | -0.106, 0.007 |
| 6 | *Bipolar Disorder* | **-0.122** | **-0.193, -0.05** | **-0.181, -0.064** |
|  | *Benzodiazepine* | 0.042 | -0.015, 0.099 | -0.007, 0.09 |

**eTable 3.** Relationship between BD and Prediction Intervals excluding the null are in bold.

| Shen Canonical Networks | Jaccard Similarity with NOI* | Mean | 95% Prediction Interval | 90% Prediction Interval |
| --- | --- | --- | --- | --- |
| FPN | 0.000 | -0.012 | -0.061, 0.038 | -0.053, 0.03 |
| Limbic | 0.000 | -0.056 | -0.125, 0.014 | -0.112, 0.003 |
| DMN | 0.118 | -0.01 | -0.066, 0.045 | -0.056, 0.035 |
| Medial Frontal | 0.000 | 0.001 | -0.044, 0.046 | -0.036, 0.039 |
| Motor | 0.030 | -0.001 | -0.049, 0.047 | -0.041, 0.04 |
| Visual Association | 0.000 | -0.021 | -0.062, 0.022 | -0.056, 0.014 |
| Visual I | 0.000 | -0.02 | -0.057, 0.018 | -0.051, 0.01 |
| Basal Ganglia | 0.306 | -0.093 | -0.178, 0.002 | **-0.166, -0.015** |
| Cerebellum | 0.021 | -0.12 | **-0.235, -0.006** | **-0.217, -0.022** |

*Jaccard Similarity

**eFigure 1:** Scanning decision-tree for BD youth. The aim of this protocol was to maximizes our chances to scan participants in different mood states.

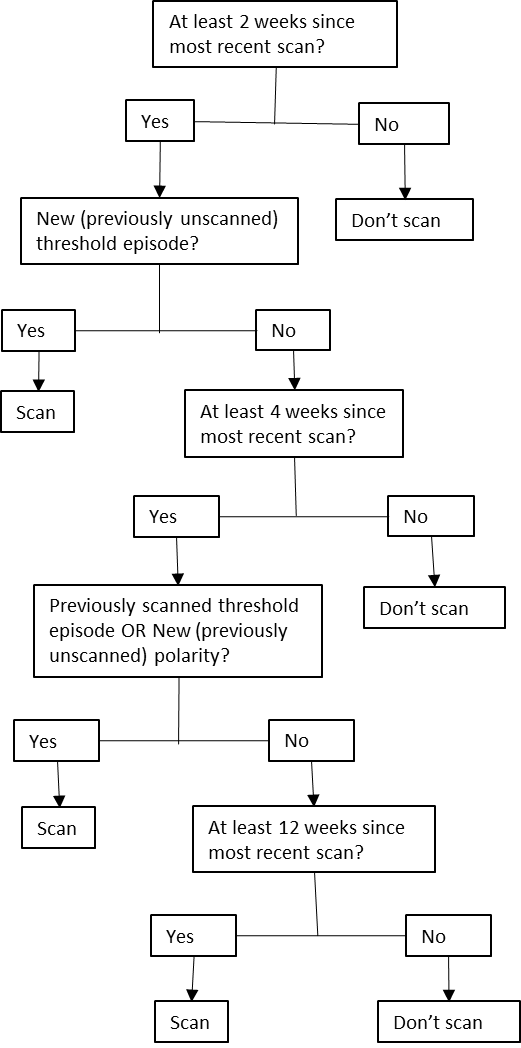

**eFigure 2:** Network of interest (NOI). This NOI was constructed based on functional connectivity patterns previously found to be associated with mania and/or depression, and overlaps extensively with regions key to reward processing. Shen parcellation labels are superimposed. The only labels not shown are parcels 126-128 (R thalamus), MNI coordinates: X=-10, Y=9, Z=-16,

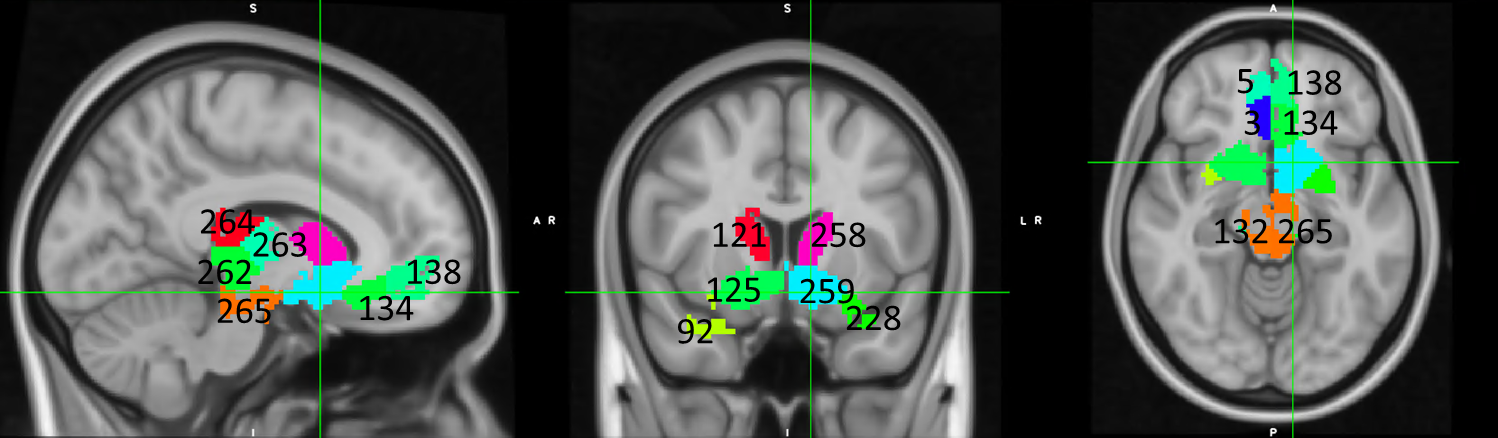

**eFigure 3:** Protocol for assessing similarity across scans in preprocessed data.

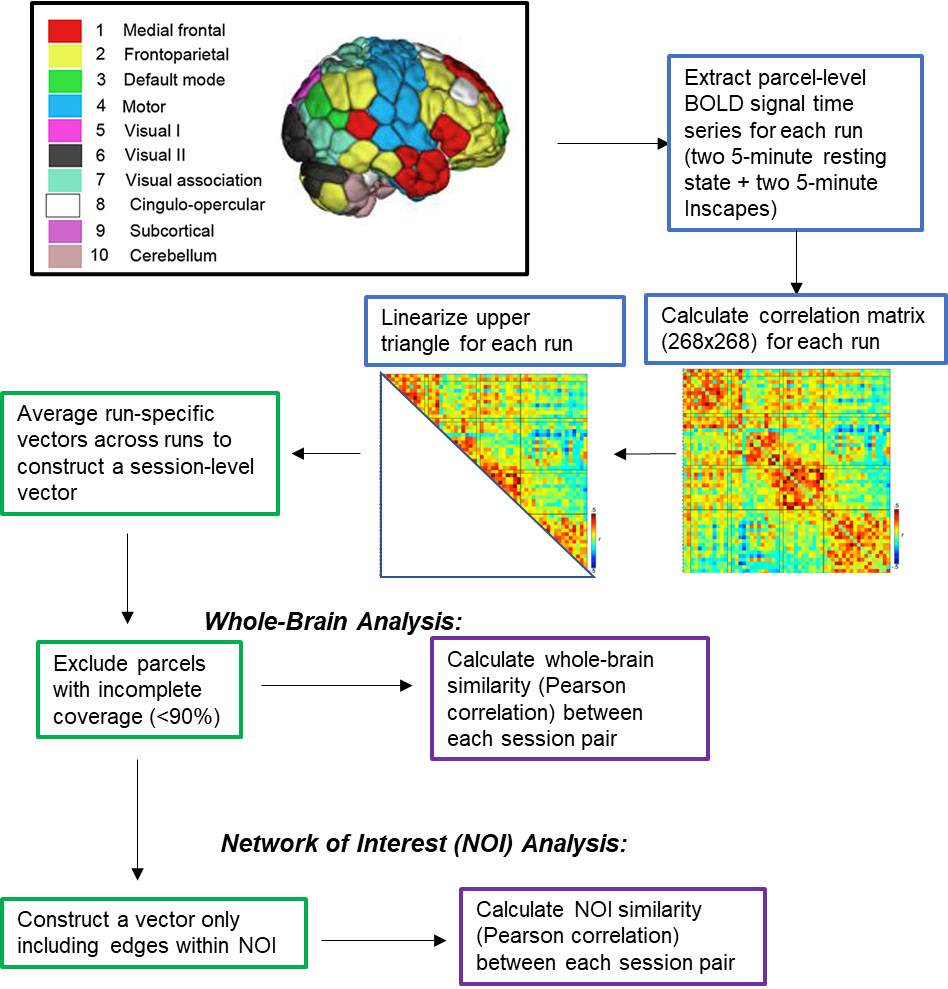

**eFigure 4:** Subject-level number of scans in different mood states

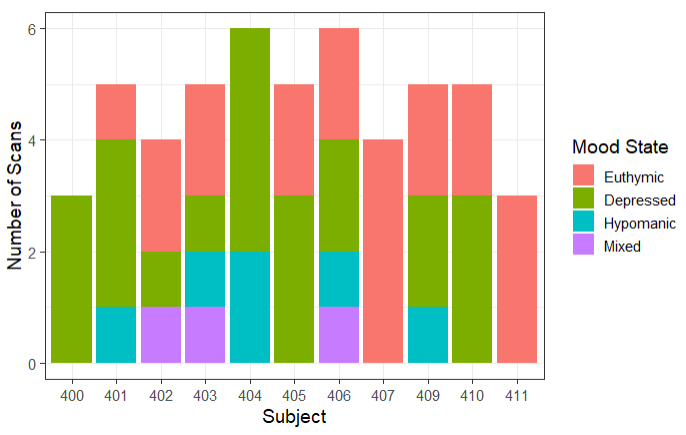

**eFigure 5:** Relationship between BD subtype and NOI stability

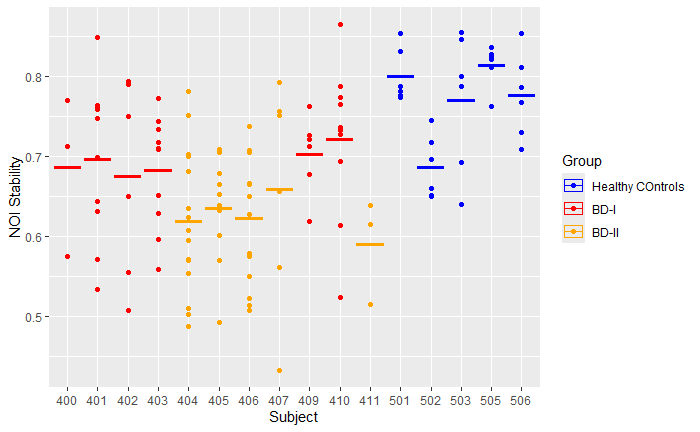

**eFigure 6:** Across-scan stability of FC in canonical Shen Networks. Differences are found primarily in the NOI (both across- and within-scan stability), but also in other subcortical networks. Cortical networks (with the highest across-scan reliability in both groups) do not show such differences.

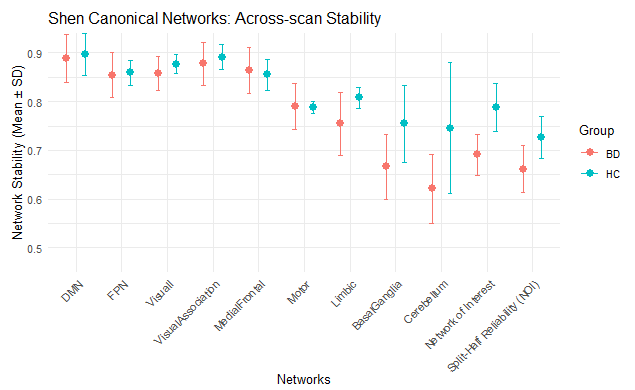
